# Face Viewing Behavior Predicts Multisensory Gain During Speech Perception

**DOI:** 10.1101/331306

**Authors:** Johannes Rennig, Kira Wegner-Clemens, Michael S Beauchamp

## Abstract

During face viewing, some individuals prefer to fixate the mouth while others fixate the eyes. Individuals who have a history of mouth fixation might have stronger associations between visual and auditory speech, resulting in improved comprehension. First, we measured eye movements during face-viewing and observed high interindividual variability in mouth fixation time. Next, we measured eye movements and comprehension during perception of noisy auditory speech with or without visual speech. When visual speech was present, participants primarily fixated the mouth, but derived substantial benefit compared to noisy auditory speech with high interindividual variability. The benefit of visual speech was predicted by the eye movements made during the initial face-viewing task, but not by eye movements during the noisy speech task. These findings suggest a link between eye movements during face viewing and audiovisual speech perception and suggest that individual histories of visual exposure shape abilities across cognitive domains.

## Introduction

When conversing, humans use visual information from the talker’s face to complement auditory information from the talker’s voice. The mouth movements made by the talker provide an independent source of information about speech content that is especially useful under circumstances in which the auditory signal is degraded, as in a noisy room. While the ability of visual speech to enhance the intelligibility of noisy auditory speech is well documented (Grant et al. 1998; Sumby and Pollack 1954; for a review see Peelle and Sommers 2015) published studies report high interindividual variability across all tested stimulus types, including consonants, words, meaningful sentences or anomalous sentences spoken in different speaking styles (Grant et al., 1998; Sommers, Tye-Murray, & Spehar, 2005; Van Engen, Phelps, Smiljanic, & Chandrasekaran, 2014; Van Engen, Xie, & Chandrasekaran, 2017); across all types of auditory noise, including multi-talker babble and speech-shaped noise (Sommers et al., 2005; Tye-Murray, Spehar, Myerson, Hale, & Sommers, 2016; Van Engen et al., 2014, 2017); and across all populations, including young and old adults (Sommers et al., 2005; Tye-Murray et al., 2016). In every study, some participants show a small benefit for visual speech while others show a large benefit.

A different axis of individual variability is found in the eye movements made by humans viewing faces. As first described by Yarbus (1967), different individuals presented with the same image make very different eye movements. Recent work has extended this finding to individual differences in face viewing. A preference to fixate the mouth or eye region of the face is found for both static and dynamic faces, is consistent across different face exemplars (Gurler, Doyle, Walker, Magnotti, & Beauchamp, 2015; Mehoudar, Arizpe, Baker, & Yovel, 2014), and is stable across testing sessions as long as 18 months apart (Mehoudar et al., 2014). Another relevant Yarbus (1967) observation was the sensitivity of eye movement behavior to task demands (Schurgin et al., 2014). During a task requiring recognition of noisy audiovisual speech, participants primarily fixate the mouth of the talker, reflecting the increased behavioral relevance of visual speech (Buchan, Paré, & Munhall, 2008; Vatikiotis-Bateson, Eigsti, Yano, & Munhall, 1998). The contributions of interindividual and inter-task differences to eye movement behavior have been integrated using Bayesian ideal observer models (Peterson & Eckstein, 2012, 2013).

Since humans view faces for thousands of hours in daily visual experience, individuals’ idiosyncratic preferences to fixate the mouth or eyes might lead to increased experience and expertise for the most-viewed face part. An individual who prefers to fixate the mouth of the face could accumulate greater expertise in decoding the visual speech information present in talker’s mouth movements and realize a greater benefit of visual speech in enhancing the intelligibility of noisy auditory speech.

To test this idea, we performed two separate experiments within the same testing session. In the first experiment, we measured participants’ preferred face viewing behavior using a stimulus in which fixating the mouth was not essential to performing the behavioral task. In the second experiment, we measured participants’ ability to understand noisy audiovisual speech, task in which information from the talker’s mouth is undeniably important. Then, we compared the results of the two experiments to determine if the two axes of variability—individual differences in face looking and individual differences in noisy audiovisual speech perception—were linked.

## Methods

### Participants, Stimuli and Task

33 native English speakers (18 female, mean age 21, range 18-32) provided written informed consent under an experimental protocol approved by the Committee for the Protection of Human Participants of the Baylor College of Medicine, Houston, TX.

#### Sample size justification

To determine the number of subjects for this study, we first collected data from 10 pilot subjects. In this initial analysis, the primary correlation of interest between speech perception and eye movements was 0.60. Recognizing this value may be inflated, we used an expected correlation of 0.50 for our power analysis. Using G*Power (version 3.1, http://www.gpower.hhu.de/en.html), we determined that we needed 30 participants to achieve 80% power. Because of the potential for data loss, we added a 10% cushion, bringing our total to 33.

Participants’ eye movements were monitored using an infrared eye tracker (Eye Link 1000 Plus, SR Research Ltd., Ottawa, Ontario, Canada) as they viewed recordings of audiovisual speech presented on a high resolution screen (Display++ LCD Monitor, 32” 1920 × 1080, 120 Hz, Cambridge Research Systems, Rochester, UK) using Matlab (The Mathworks, Inc., Natick, MA, USA) with the Psychophysics Toolbox extensions (Brainard, 1997; Pelli, 1997). Stable observed head positioning was ensured with a chin rest placed 90 cm from the display. Auditory stimuli were presented through speakers on either side of the screen.

Participants viewed two kinds of speech stimuli. In the first experiment (Figure 1A), the stimuli consisted of audiovisual recordings of single syllables without any added noise (clear syllables). Each trial began with a fixation crosshairs presented outside of the location of where the face would appear in order to simulate natural viewing conditions in which faces do rarely appear at the center of gaze (Gurler et al., 2015). As soon as the audiovisual speech video began playing, the fixation crosshairs disappeared and participants were free to fixate anywhere on the screen. After the speech video ended (2 seconds duration) participants reported the identity of the syllable with a button press. Each participant viewed 270 syllable trials (divided into two runs): 20 repetitions × 4 talkers × 3 audiovisual syllables (2 congruent: AbaVba, AgaVga; 1 incongruent: AbaVga) and 10 repetitions × 1 talker × 3 audiovisual congruent syllables (AbaVba, AgaVga, AdaVda), all randomly interleaved; only the congruent syllable data was analyzed. Participants identified the congruent syllables (AbaVba, AdaVda and AgaVga) with 96% accuracy (SD 7%, range 63’ 100%).

**Figure 1.**
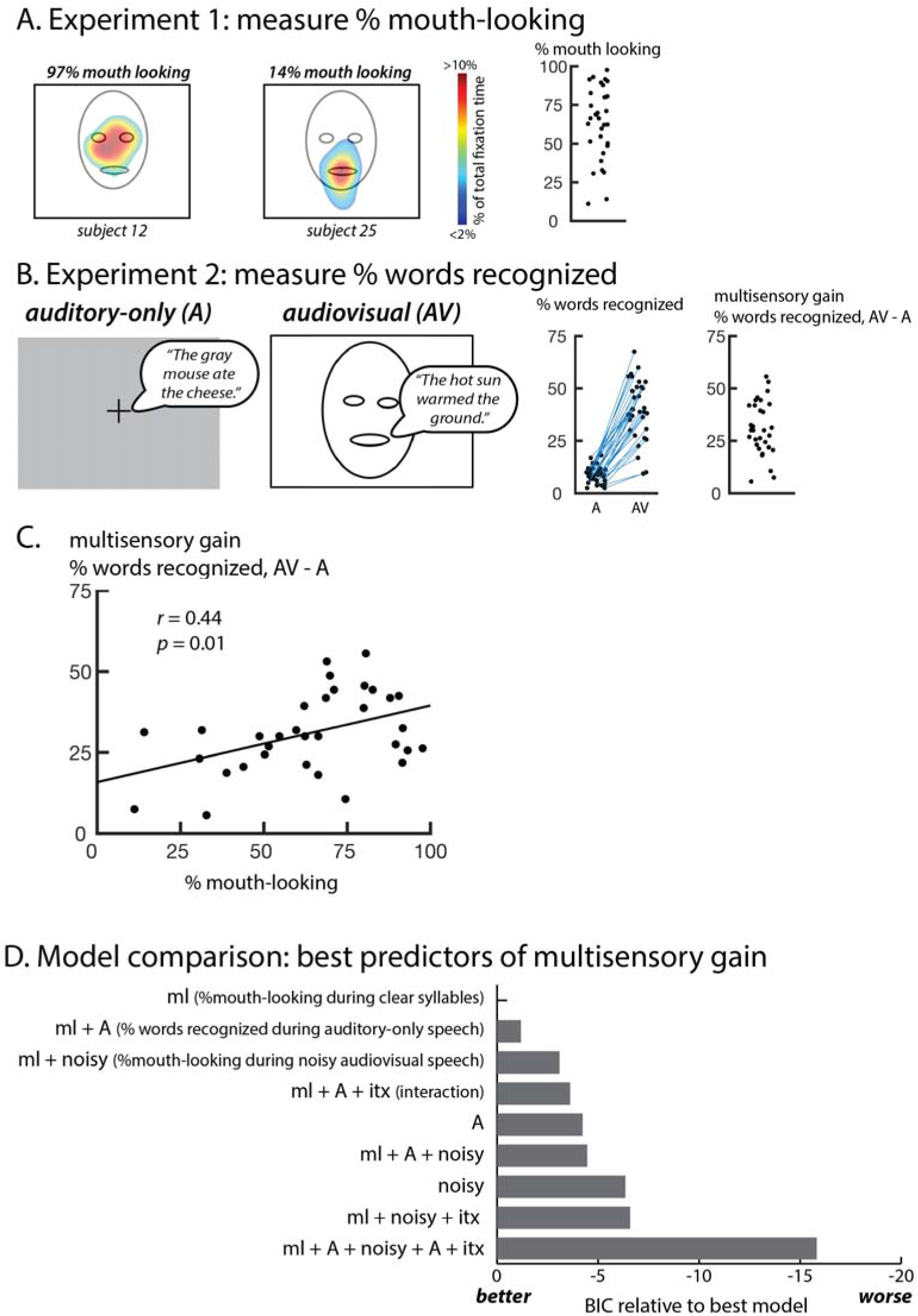
Face Viewing Behavior Predicts Multisensory Gain During Speech Perception. **A.** In the first experiment, eye movements were measured during face-viewing. Colors (overlaid on a schematic still frame from the stimulus video) show the time spent fixating each location in the display as a percentage of total fixation time for two sample participants. The fixation maps were converted to a single number, the percentage of time fixating the mouth region of the face (defined as the lower half of the stimulus display). The plot shows this value for all participants, one symbol per participant. **B.**In the second experiment, participants reported the words in noisy sentences presented without a video of the talker’s face (auditory-only, A) or with the video (audiovisual, AV). Speech bubble shows the auditory component of the stimulus. The left plot shows the percentage of key words recognized as a fraction of total words in each condition (two symbols for each participant connected by a blue line). The right plot shows multisensory gain, calculated as the difference between the two conditions (one symbol per participant). **C.** Correlation across participants between mouth-looking time (plot in **A**) and multisensory gain (right plot in **B**) with one symbol per participant. **D.** In order to determine the factors influencing multisensory gain, models were constructed that included all measured experimental variables. The models were compared using the Bayesian Information Criterion (BIC) which adjusts the total variance explained by the number of predictors, penalizing overfitting. Models are ordered from best to worst.

In the second experiment (Figure 1B), the stimuli consisted of auditory sentences recorded from a single male talker with added auditory noise (pink noise, signal-to-noise ratio 16 dB) presented either alone (auditory-only) or paired with a video recording (audiovisual) (Van Engen et al. 2017). Each trial began with a fixation crosshairs presented outside of the location of where the face would appear in audiovisual trials. During auditory-only trials, the fixation crosshair shifted to the center of the screen when auditory playback began. In audiovisual trials, the fixation crosshair disappeared when the video began. After the sentence ended (3 seconds duration) participants repeated the sentence. Responses were scored for number of correct keywords (*e.g.* “The **hot sun warmed** the **ground**,” keywords in bold). Each participant viewed 80 sentence trials (divided into two runs): 40 auditory-only and 40 audiovisual, interleaved with no sentences repeated. For each participant, the total number of keywords recognized was divided by the total number of keywords to generate a “percentage words recognized” score for the auditory-only and audiovisual conditions.

### Eye tracking and analysis

Eye tracking was performed with a sampling rate of 500 Hz. Before each run of each task, a 9-target array was presented for eye-tracker calibration and validation. Three times within each run, participants fixated a centrally presented crosshair. The difference between the measured eye position during these epochs and the screen center was applied to correct the eye tracking data in the preceding stimulus epoch. Two regions of interest (ROIs) were defined for each video, consisting of an upper face (eye) ROI and a lower face (mouth) ROI (Figure 1A). Blinks and saccades were excluded from the analysis and the percentage of fixation time spent within each ROI was calculated.

## Results

In the first experiment, participants identified the syllable spoken by an audiovisual talker. All participants were at ceiling, similar to a previous study in which auditory-only versions of the same stimuli were presented (mean of 98% accuracy, range 88 to 100%, in the present study *vs.* mean of 97% in Mallick et al. 2015), demonstrating that distinguishing single syllables is an easy task that does not require visual speech information. While participants were at ceiling accuracy, there was high variability in eye fixation behavior. Some participants spent as little as 11% of total fixation time fixating the mouth of the talker, while others spent as much as 98%, with a mean of 64% (Figure 1A).

In the second experiment, participants listened to noisy auditory sentences presented with a fixation crosshair (auditory-only) or paired with a video of the talker’s face (audiovisual) (Figure 1B). A high level of auditory noise was used (−16 dB) with the result that in the auditory-only condition participants recognized only a few words (mean 9%, range: 3 to 18%). In the audiovisual condition, participants recognized many more words (mean 40%, range: 9 to 68%; paired *t*-test *t*(_32_) = 14.4, *p* = 1 × 10^−15^), demonstrating the benefit of visual speech in enhancing the intelligibility of noisy auditory speech.

Every single participant showed improved performance between the auditory-only and audiovisual speech conditions. The amount of improvement, referred to as multisensory gain (calculated as the % words recognized during audiovisual noisy speech – % words recognized during auditory-only noisy speech) quantifies the benefit provided by viewing the talker’s face. The mean multisensory gain was 31% but there was a high degree of variability. Some participants improved as little as 6% while others improved as much as 56%.

We observed high interparticipant variability in two measures: eye movements during viewing clear syllables and multisensory gain for noisy audiovisual speech. To determine if there was a relationship between these measures, we plotted them against each other (Figure 1C). Participants who spent more time fixating the mouth during viewing of clear syllables had higher multisensory gain (Figure 1C; *r* = 0.44, *p* = 0.010; effect size: 0.66). The slope of this curve was 0.46: for each 10% extra time a participant spent observing the mouth during clear syllables, their multisensory gain increased by 4.6%.

One possible explanation for these results is that mouth-looking behavior during presentation of clear syllables simply reflects mouth looking during noisy speech perception. However, in contrast to the variability in eye movements during viewing of clear syllables, during viewing of noisy sentences all participants primarily fixated the mouth (Figure 2A; mean of 93% of total fixation time, range from 72% to 100%). Reflecting this lack of variability, there was no significant correlation between the percent total time spent fixating the mouth during noisy sentences and multisensory gain (*r* = 0.05, *p* = 0.773).

**Figure 2.**
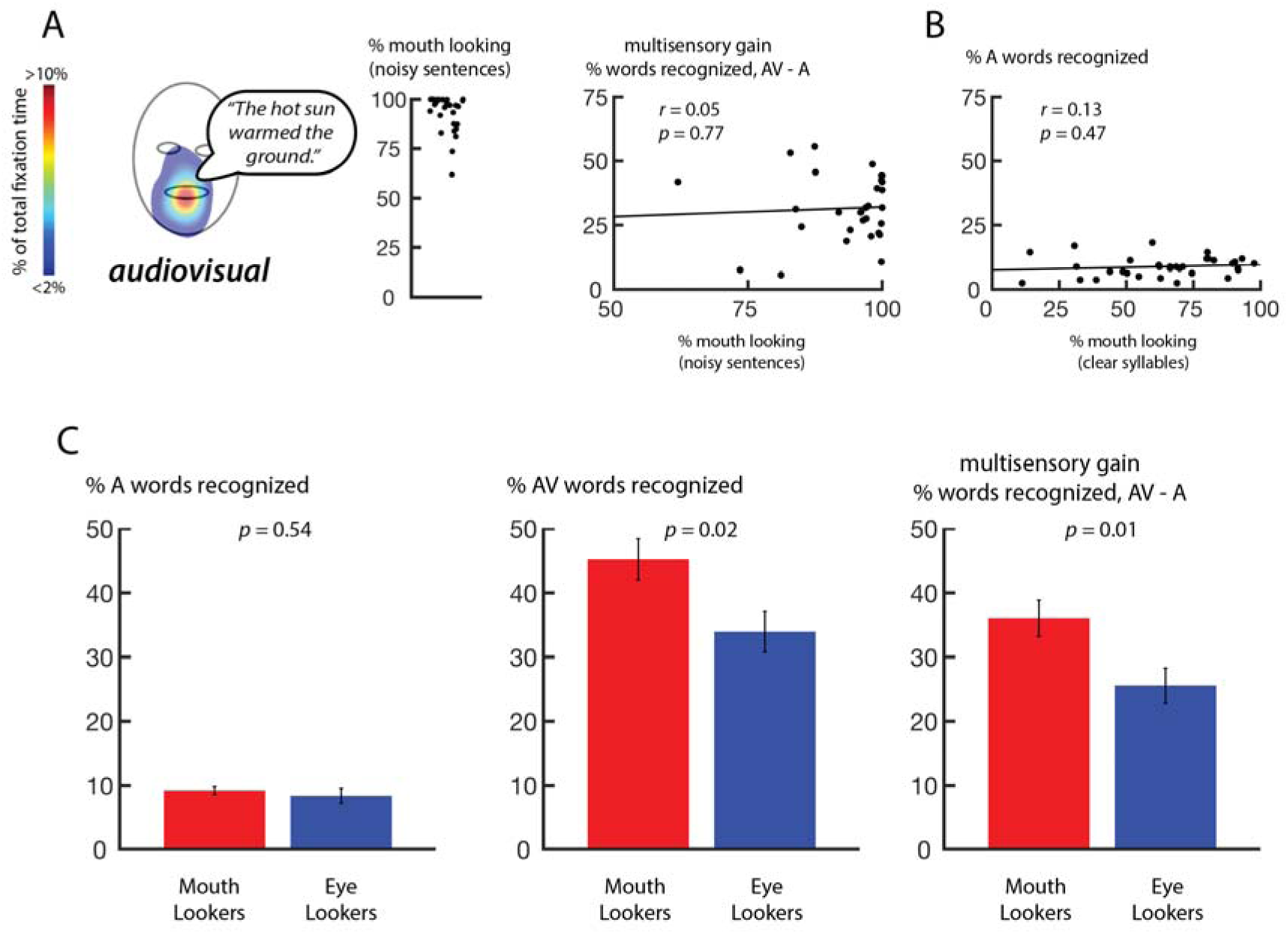
**A.** Eye movement measurements from the second experiment, perception of noisy audiovisual sentences. Colors (overlaid on a still frame from the stimulus video) show the time spent fixating each location in the display as a percentage of total fixation time, averaged across participants (speech bubble shows auditory stimulus). Left plot shows the percentage of time fixating the mouth region of the face, one symbol per participant. Right shows the correlation between this value and multisensory gain (right plot in **Figure 1B**). **B.** Correlation between mouth-looking (**Figure 1A**) and noisy auditory word recognition (left plot in **Figure 1B**). **C.** A median split was applied to the eye movement data from experiment 1 to classift participants as “mouth lookers” or “eye lookers”. For this grouping, bar graphs show percentage of noisy auditory words recognized, percentage of noisy audiovisual words recognized, and the multisensory gain (difference between audiovisual and auditory). Error bars show the standard error of the mean.

Another possible explanation of these results is that mouth-looking behavior during presentation of clear syllables reflects ability to understand noisy auditory-only speech. However, we observed no correlation between mouth-looking and auditory-only recognition (Figure 2B; *r* = 0.13, *p* = 0.473).

To integrate these findings, we constructed nine different models that used all of our measured variables and their interactions to predict multisensory gain. Bayesian model comparison was used to determine the model that best predicted performance of individual participants. Overfitting with excess parameters was penalized using the Bayesian information criterion (BIC). As shown in Figure 1E, the winning model used only a single variable—mouth-looking during viewing of clear syllables—to predict multisensory gain. The next-best model (three times less likely than the winning model given the observed data) included an additional variable, auditory-only noisy speech recognition. To understand the contribution of this variable, we plotted it against multisensory gain but did not find a significant correlation between A-only performance and multisensory gain (*r* = 0.29, *p* = 0.103).

For comparison with previous studies (*e.g.* Gurler et al. 2015), we applied a median split to classify participants into one of two categories: those who most often fixated the mouth of the talker during viewing of clear syllables (mouth lookers) and those who less often fixated the mouth preferring the eyes of the talker (eye lookers). We constructed a linear mixed-effects model with participant as a random factor and two fixed effects for stimulus type (auditory-only *vs.* audiovisual) and group (mouth-lookers *vs.* eye-lookers). The dependent measure was the percentage of words recognized, with the baseline condition consisting of the factor levels “auditory-only” and “mouth-lookers.” (lme4 package in R; Bates et al. 2015). There was a large main effect of condition reflecting enhanced recognition for audiovisual speech (parameter estimate for difference in words recognized: 25%; *F*(1,33) = 258.80, *p* = 2 × 10^−16^) and a significant effect of group reflecting improved performance by mouth-lookers (+1%; *F*_(1,33)_ = 5.46, *p* = 0.026). Critically, there was a significant interaction between condition and group, driven by greater multisensory gain for mouth-lookers (+10%; *F*(_1,33_) = 7.44, *p* = 0.010; Figure 2C).

## Discussion

Yarbus (1967) first demonstrated that individuals make varied eye movements when confronted by a visual image and that this variability has two components, an inter-task component and an inter-individual component. The inter-task component is results from humans modifying their eye movement behavior based on task demands. For instance, when asked to identify joy, an emotion that is primarily represented in the mouth region of the face, humans are more likely to fixate the mouth (Schurgin et al., 2014). Similarly, when perceiving noisy auditory speech, humans are more likely to fixate the mouth than when perceiving clear auditory speech, because the noisy speech task benefits from visual speech information (Buchan et al., 2008; Vatikiotis-Bateson et al., 1998). We replicated this finding in our study, with the average time spent fixating the mouth increasing from 64% during the clear speech task of experiment 1 to 93% in the noisy speech task of experiment 2.

The inter-individual component of eye movement variability is less well understood. Recent studies have shown that different individuals have idiosyncratic preferences in how they view faces (Gurler et al., 2015; Mehoudar et al., 2014; Peterson & Eckstein, 2012, 2013). Even when performing the identical task, some individuals prefer to fixate the mouth of the talker while others fixate the eyes, a preference that is unchanged when tested up to 18 months apart (Mehoudar et al., 2014; Peterson & Eckstein, 2012, 2013). Interindividual differences in face preference has been shown for both static faces (Mehoudar et al., 2014; Peterson & Eckstein, 2012, 2013) and dynamic talking faces (Gurler et al., 2015) and the contributions of interindividual and inter-task differences to eye movement behavior have been integrated using Bayesian ideal observer models (Peterson & Eckstein, 2012, 2013). In our first experiment, we first measured participants’ face movement behavior using an undemanding task (recognizing clear speech in a quiet room). Since this task can be performed with near perfect accuracy even without visual speech (Mallick et al., 2015), face viewing behavior was driven by participants’ internal preferences rather than by task demands. Consistent with previous studies, we observed substantial interindividual variability, with mouth fixation time ranging from 11% to 98% of total fixation time.

Another poorly understood axis of individual variability is the perceptual benefit provided by viewing a talker’s face. While the large benefit of seeing the face in understanding noisy speech is incontrovertible (Grant et al. 1998; Sumby and Pollack 1954; for a review see Peelle and Sommers 2015) there is no explanation in the literature for the high interindividual variability in this benefit that is observed across all published experiments (Grant et al., 1998; Sommers et al., 2005; Tye-Murray et al., 2016; Van Engen et al., 2014, 2017). Consistent with these reports, we observed large individual variability, with audiovisual gain ranging from 9 – 68% across participants.

We hypothesized that individuals with a preference for fixating the mouth of a viewed face would have an advantage in processing visual speech. To test this hypothesis, we correlated the face looking behavior from experiment 1 with noisy speech identification from experiment 2. We observed a significant correlation between individual difference measures of viewing behavior and audiovisual gain. This finding could not be explained by eye movement behavior in experiment 2, in which the task requirements led all participants to fixate the talker’s mouth. Instead, participants who fixated the mouth when it was not important (during the initial face-viewing task) received more benefit from fixating the mouth when it was important (during the noisy speech task). Conversely, even fixating the mouth when it was important (during the noisy speech task) provided less benefit to participants who did not fixate the mouth when it was not important. These observations were confirmed with Bayesian model comparison. The best-performing model predicted noisy speech perception only using eye movement behavior during face-viewing.

A natural question for future studies is whether eye lookers have an advantage in other tasks as compensation for their worse performance in speech perception. The eye region of the face is important for face memory (Peterson & Eckstein, 2012, 2013). Therefore, we speculate that eye viewers, with more experience at the eye region of the face, could be better at face memory.

Faces are one of the most important classes of visual stimuli, as evidenced by the many brain areas that have evolved in primates to process face information (Grill-Spector, Knouf, & Kanwisher, 2004; Haxby, Hoffman, & Gobbini, 2002; Kanwisher & Yovel, 2006) with regions specialized for extracting the different types of information available in the eye and mouth regions of the face (Pelphrey, Morris, Michelich, Allison, & McCarthy, 2005; Zhu & Beauchamp, 2017). The quality of the information available to the brain about different facial features is determined by gaze location. Since the retina has a region of very high acuity in the fovea, fixating a specific region of the face provides higher quality information about the relevant stimulus features. This would provide a potential neural mechanism for accumulating expertise about different facial features that would be different across individuals, depending on their preferred fixation location when viewing a face. An important concept in vision science is that the visual system is tuned by exposure to natural visual scenes. Our work extends this concept by showing that individual differences in patterns of visual exposure, shaped by eye movements, can lead to substantial individual differences in perception.

## Acknowledgments

This work was supported by the National Institutes of Health (R01NS065395 to M.S.B) and the Deutsche Forschungsgemeinschaft (RE 3693/1-1 to J.R.). We acknowledge the Core for Advanced MRI at Baylor College of Medicine. We thank Bharath Chandrasekaran for providing the stimuli for the noisy sentence experiment.

## Author contributions

MSB and JR developed the study concept and study design. Data collection was performed by KWC. JR and KWC conducted data analysis. MSB and JR wrote the paper.

